# Dietary reconstructions of Magdalenian canids from SW-Germany do not indicate that the area was a centre of early European wolf domestication

**DOI:** 10.1101/2023.11.27.568675

**Authors:** Paul D. Bons, Catherine C. Bauer, Lydia J. Papkalla

**Affiliations:** Department of Geosciences, Tübingen University, Schnarrenbergstrasse 94-96, 72076 Tübingen, Germany

## Abstract

In their paper *‘A refined proposal for the origin of dogs: the case study of Gnirshöhle, a Magdalenian cave site’*, Baumann and colleagues^[1]^ claim that their data *‘support the hypothesis that the Hegau Jura was a potential center of early European wolf domestication’*, and that *‘such a scenario becomes plausible considering a close proximity of canids and humans thereby introducing a controlled, or at least a restrictive diet’*. The study focusses on fossil remains of *‘large canids’* from the Gnirshöhle cave site in SW Germany. Morphometric data on only one specimen, GN-999, as well as collagen δ^15^N and δ^13^C isotopic data and mitochondrial DNA analyses on the Gnirshöhle specimens and a comparative sample were used to conclude that the Gnirshöhle specimens shed light on the *‘origin of dogs’* as purported by the title of the paper. Here we argue that the paper is fundamentally flawed and excluded available relevant data.

## Cluster analysis and trophic niches

Both the morphometrics and the mitochondrial DNA were found to be inconclusive to assign the Gnirshöhle specimens to either dog or wolf. The authors therefore heavily rely on the ‘trophic niches’ concept based on bulk collagen δ^13^C and δ^15^N analyses (their Fig. 3). The authors derive three ‘trophic niches’, one of which comprises six Gnirshöhle specimens together with the three Kesslerloch dogs and one red fox, also from Kesslerloch. This ‘trophic niche’ is assigned to ‘dogs’ (as the dog drawn in their Fig. 3 graphically suggests). No arguments are provided for this assignment to a group in which six out of ten specimens are ‘large canids’ of undetermined status (to be discussed below) and one is a red fox. In a next step dietary reconstructions are provided that show clear differences between the diets of the three ‘niches’ (their Fig 3C), which is then finally used to make a case for the origin of dogs, although no evidence is presented that any of the Gnirshöhle specimens is a dog. Apart from this, there are a number of critical issues with the application of the trophic niche concept and the dietary reconstruction.

The basic idea of the dietary reconstruction^[2,3,4,5,6,7]^ is that the isotopic composition of the diet of an animal is reflected by the isotopic composition of that animal’s tissues^[2,3,8]^. Grazers will record the isotopic signature of the vegetation they consume and predators in turn that of the grazers. Trophic niches are defined as groups of animals that shared a common diet and thus have similar δ^13^C and δ^15^N isotopic signatures in their tissues. In isospace (a biplot of δ^15^N *versus* δ^13^C) the isotopic data of a group or niche would cluster together. This, however, does not mean that any cluster in isospace is automatically a trophic niche^[9]^. We here do not wish to challenge the powerful methods of dietary reconstructions and the useful concept of niches^[10,11]^, but only the way it is implemented by Baumann and colleagues^[1]^.

Baumann and colleagues^[1]^ define three ‘geometric’ isospace clusters (their Fig. 3a), and without further evidence equate these with ‘trophic niches’. With ‘geometric’ we mean that these clusters are purely based on the position of the specimens in isospace and do not take any other biological or ecological information into consideration. They fail to provide details of their cluster analysis. All that is reported is that the software JMP 14 was used. From this we may assume that JMP’s default Ward’s method with standardisation of data was used. There is no discussion on why this is the preferred or appropriate method. Furthermore, there is no discussion on the criteria to choose three clusters, and not, for example, two (low and high δ^15^N) or more. In order to equate clusters with trophic niches one should first set criteria to define under which conditions a cluster in isospace would constitute a trophic niche. Such criteria are not provided, and it therefore remains unknown how statistically significant the three ‘trophic niches’ are, and even less if they have any ecological meaning or significance.

Although the isotopic signature of the tissues of a predator reflects that of its prey, it is not identical, but shifted due to fractionation by a trophic enrichment factor or TEF (ΔX)^[8,12,13,14]^: δXpredator = δX_prey_ + ΔX, where X is the isotope under consideration, here ^13^C or ^15^N. The TEF depends on a number of factors, such as species, age, and health of the individual, or even the isotopic value of the source^[14,15,16]^. Baumann and colleagues^[1]^ use Δ^13^C = 1.1 ± 1.1‰ (one δ) and Δ^15^N = 3.2 ± 1.8‰. Individuals with an identical diet may record a spread in isotopic signatures in the order of the standard deviation. Alternatively, individuals with a diet that differs by 1-2 ‰ may potentially record identical isotope values in their tissues. This is not a major concern when the sample size is large enough. However, here we have clusters of only five to ten individuals that are separated by less than one standard deviation of the TEFs.

A reasonable null-hypothesis that should have been considered would be that all specimens had the same diet and that the spread in individual isotopic signatures is the result of the variability in which individuals record the isotopic signatures of their prey. In that hypothetical case we would expect the Magdalenian canids to show δ^13^C values with a standard deviation *σ*_C,sample_≤1.1 ‰ and δ^15^N values with *σ*_N,sample_≤1.8 ‰. In fact, the twenty specimens have *σ*_C,sample_=0.5 ‰ in their δ^13^C values, and *σ*_N,sample_=1.2 ‰ for δ^15^N, which is well within what would be expected if all specimens had the same diet. Proper arguments should have been provided by Bauman and colleagues^[1]^ to argue for a ‘restrictive’ diet of a subgroup that is supposedly related to the domestication of dogs.

The above is based on statistical considerations only. It may also be useful to consider what the known spread in measured isotopic signatures of wolves can be. Different studies have determined different TEFs for the different tissues of wolves^[17,18,19,20]^, but here we consider collagen for the Magdalenian specimens and hairs for modern wolves. A classical study on wolves from coastal British Columbia^[21]^ analysed the isotopic signatures of hairs of wolves in three distinct regions (mainland, inner islands adjacent to the coast, outer islands). Of these three regions, the mainland one is probably the most comparable with Magdalenian SW Germany where marine mammals were absent, but some salmon or other fish could potentially have been consumed. For the British Columbia study, it was known of each wolf to which pack it belonged. It can be assumed that members of one pack had approximately the same diet. We see (Fig. 1a) that the variability in isotopic signature of the 21 mainland wolves is comparable to that of the 20 specimens of the study of Baumann and colleagues^[1]^. It should be noted here that the British Columbia data were collected in a four-year period in a well-defined area and with well-constrained prey groups. It should be borne in mind that the isotopic signature of hairs is accumulated in a short time, compared to that of bone collagen, which may increase isotopic variability^[22,23,24]^. However, the canid and prey specimens in the study of Baumann and colleagues^[1]^ may also have an increased variability because they come from France and SW Germany and cover a period of thousands of years in the Magdalenian, which lies in the transition between the Last Glacial Maximum and the significantly warmer beginning of the Holocene. Variations in ecosystems and baseline δ^13^C and δ^15^N are expected because of this^[25,26]^. figure 1a thus supports the abovementioned null-hypothesis that the Magdalenian specimens cannot be divided into meaningful ‘niches’ based on the available isotopic data only.

**Figure 1.**
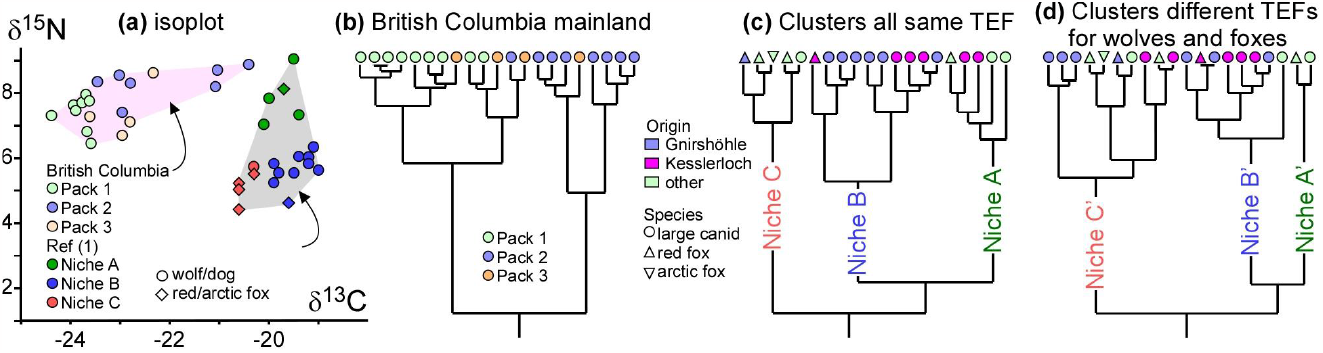
(a) Isoplot of mainland wolf packs (measured on hairs) in British Columbia^[21,29]^ and of the canids in the study of Baumann and colleagues1 (measured on collagen). Colours represent individual packs (British Columbia) or ‘trophic niches’ (Magdalenian specimens^[1]^). (b) The clusters obtained from a cluster analysis of the 21 British Columbia wolves do not coincide with the known three wolf packs. (c) Cluster analysis of the Magdalenian specimens, assuming the same TEF for all specimens, shows a comparable clustering in terms of distances as for (b), although the individual clusters include specimens of different species and origin. (d) Cluster analysis of the same Magdalenian specimens considering different TEFs for large canids^[17]^ and for foxes^[31]^. This results in very different clusters. Cluster analyses were performed using Ward’s method with normalised data in JMP 16.0.

Baumann and colleagues^[1]^ applied a cluster analysis to define clusters as ‘trophic niches’. We applied the same to the 21 British Columbia Mainland wolves (Fig. 1b). We used Ward’s method with normalised data implemented in JMP 16.0. The clusters do not fully coincide with the three packs, showing that a purely ‘geometric’ or ‘blind’ cluster analysis is of little value here. A similar cluster analysis was applied to the measured isotopic values of the Magdalenian specimens (Fig. 1c) and resulted in the same three clusters as Baumann and colleagues1 defined and which they equated to ‘trophic niches’. The clusters are only based on isotopic values and do not correlate with species nor with sample location (Fig. 1c). The assumption that the three Magdalenian clusters are ‘trophic niches’ is a major leap of faith, considering that they represent three species from a variety of origins and ages that may differ by thousands of years.

### Dietary reconstructions

Baumann and colleagues^[1]^ provide dietary reconstructions for their three clusters, which are assumed to represent ‘trophic niches’. The dietary reconstructions are based on the premise that the isotopic signature of a predator is the weighted average of the isotopic signatures of its prey plus the shift due to the TEF^[2,3,4,8]^. A dietary reconstruction attempts to determine the weights, i.e., the proportions of the prey species in the diet of a predator. The inversion of the weighted average equation is far from trivial as isotopic signatures of both source (prey) and consumer (predator) vary, and the TEFs also carry an uncertainty^[13,15,16,18,27]^. Baumann and colleagues^[1]^ used the R-based Bayesian modelling software MixSIAR^[28,29]^ to determine the dietary mixes of the three niches.

Baumann and colleagues^[1]^ used isotopic data of a variety of potential prey species of Magdalenian age from France and Germany and grouped them in three groups. Although Philips and colleagues^[30]^ write ‘*common sense, such as graphing data before analyzing, is essential to maximize usefulness of these* [isotopic mixing models] *tools*’, Baumann and colleagues^[1]^ only provided the data in tables, so we here show them in isoplots (Fig. 2). The statistics of the ‘megaherbivores’ group are poor as it consists of only three specimens. Woolly rhinoceros specimens were assigned to ‘ungulates’, without further explanation. The ‘ungulates’ group is strongly dominated by reindeer that constitute 44 out of 69 specimens. Reindeer tend to have higher δ^13^C values than other ungulates. The convex hulls of ‘ungulates’ and ‘small game’ (n=21) overlap significantly, with the overlap roughly covering the space occupied by the non-reindeer ungulates.

**Figure 2.**
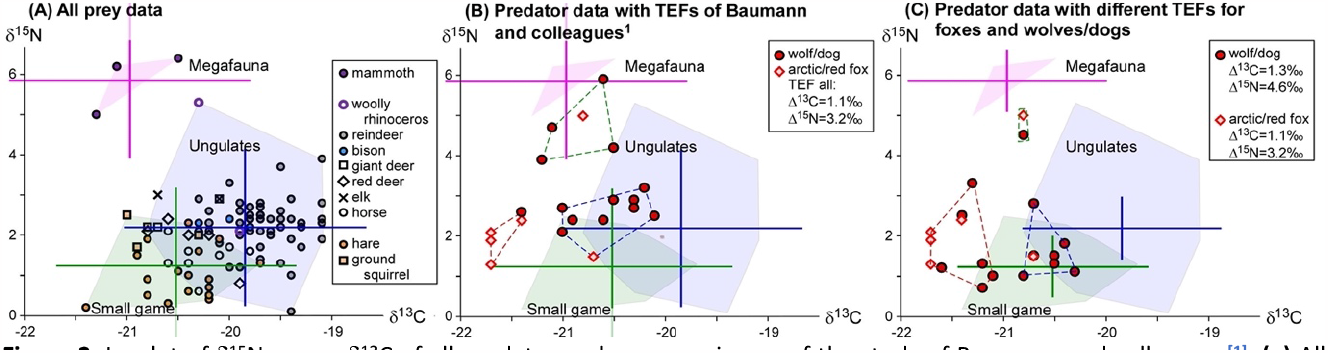
Isoplot of δ^15^N versus δ^13^C of all predator and prey specimens of the study of Baumann and colleagues^[1]^. (a) All prey data, also showing convex hulls of the three prey groups. (b) All canid data, offset according to the TEFs for foxes of Krajcarz and colleagues^[31]^, as used by Bauman and colleagues^[1]^. (c) Same canid data, but data for wolves/dogs are now offset according to the TEFs that were determined for wolves by Fox-Dobbs and colleagues^[17]^. Clustering now results in different ‘niches’ (Fig. 1d), changing the compositions, shapes and positions of the three ‘niches’ significantly. The low-δ^13^C ‘niche C’ is nevertheless always almost completely outside the convex hull of all prey data. Means are shown for the prey groups. Error bars are the combined standard deviations of the variation in sources and TEFs from Krajcarz and colleagues^[31]^ in (a) and (b) and from Fox-Dobbs and colleagues^[17]^ in (c).

**Figure 3.**
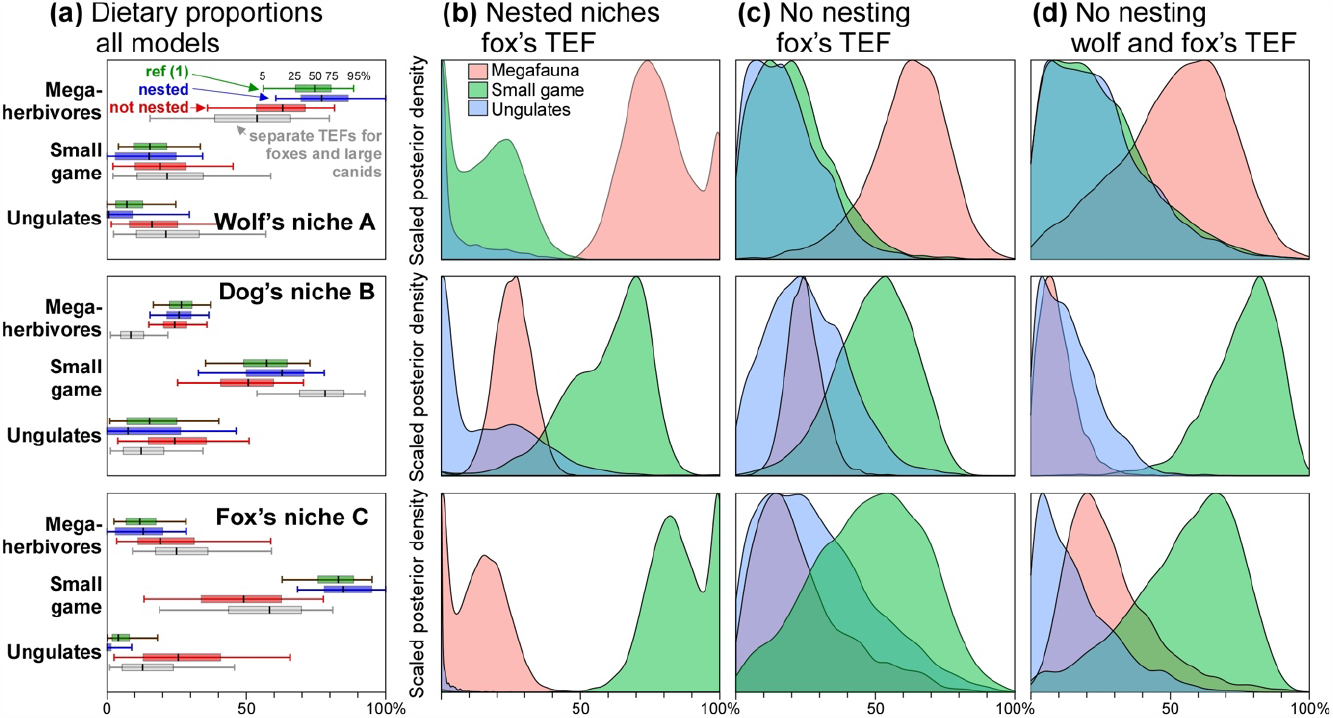
Dietary reconstructions for the three ‘trophic niches’ of Baumann and colleagues^[1]^. (a) Estimated dietary proportions for the three ‘trophic niches’ according to Baumann and colleagues^[1]^ (green), our own MixSIAR model runs with (blue) and without (red) nesting for the clusters as defined by Baumann and colleagues^[1]^, and finally our model run (grey) for the clusters shown in Figs. 1d and 2c. (b) Hierarchical (nested) reconstruction resulting in source contributions close to those reported by Baumann and colleagues^[1]^. (c) Non-hierarchical reconstruction where the dietary mix of each ‘trophic niche’ is modelled individually without consideration of the isotopic signatures of the sample as a whole. This significantly changes the dietary reconstructions, and especially increases the contribution of the ‘ungulates’ in the modelled diet of all ‘trophic niches’. (d) Non-hierarchical reconstruction for the diet of the three clusters based on separate TEFs for foxes and large canids (Fig. 2c).

The advantage of MixSIAR is that its Bayesian solver includes the variability in isotopic signature of the dietary sources. It does not return a single best-fit value for the prey mix of the predator, but the probability of the contribution of one prey species or group in the total diet of a predator. MixSIAR requires the TEFs and their standard deviations as input. Here the authors use as TEFs Δ^13^C = 1.1 ± 1.1‰ and Δ^15^N = 3.2 ± 1.8‰, which is taken from a study on modern foxes by Krajcarz and colleagues^[31]^. An earlier study from the same laboratory on the possible early domestication of wolves^[32]^ used Δ^13^C = 1.1 ± 0.2 ‰ and a Δ^15^N = 3.8 ± 1.8 ‰ for wolves. Fox-Dobbs and colleagues^[17]^ studied collagen TEFs of wild wolves from the Isle Royale National Park (USA) and obtained a Δ^13^C = 1.3 ± 0.6 ‰, and a Δ^15^N = 4.6 ± 0.7 ‰. The uncertainty in TEFs is probably the most critical factor in dietary reconstructions^[15,16,18,23,27,30]^, as small changes in the TEF can dramatically change the results, especially when the range in dietary sources is small, as is the case here for the overlapping ‘reindeer’ and ‘small game’ convex hulls. The authors do not discuss nor explain why they chose to apply TEFs for foxes to both large canids and foxes, although foxes are not really topic of the study. It is actually unclear why foxes are included at all in a study on the difference between Magdalenian wolves and dogs.

Any dietary reconstruction using isotopes should show the consumers and sources in an isoplot^[29,30]^, but Baumann and colleagues^[1]^ did not do so. Such a plot allows a visual assessment of the plausibility of the analysis and we therefore here provide it for the data of Baumann and colleagues^[1]^ (Fig. 2) When the raw data are plotted, consumers and sources will be offset by the TEFs. The plot should thus instead show the data after correcting for these TEFs (Figs 2a-b). We do so using the TEFs for foxes^[31]^ for both foxes and large canids in Fig. 2b, and using the TEFs for wolves^[17]^ for the large canids (Fig. 2c)

A striking feature of Fig. 2 is that the canids plot in a quite low δ^13^C range and several (the whole of ‘trophic niche’ C) plot outside the triangular convex hull of the source means, and also outside the convex hull of all individual prey specimens if we exclude one single outlier hare. This normally indicates that either the TEFs are wrong or one or more dietary sources are missing (here with low δ^13^C). However, MixSiAR essentially provides a best-fit solution for the data that are provided. The manual^[29]^ therefore warns that MixSIAR “will always find a solution *even if it is nonsensical*”. Not surprisingly, MixSIAR calculated a high percentage of ‘small game’ in the diets of all ‘trophic niches’ and relatively small percentages of ‘ungulates’ (their Fig. 3c).

Unfortunately, Baumann and colleagues^[1]^ did not provide essential information on their dietary reconstruction. They merely state that they used the Bayesian statistic model MixSIAR. We attempted to replicate (Fig. 3a,b) the dietary reconstructions reported by Baumann and colleagues1 (Suppl. Information). The closest match was obtained using the example script “mixsiar_script_wolves_normal.R” that accompanies a standard MixSIAR installation and is explained in the user manual^[29]^. This script was originally written for a hierarchical mixing model^[21]^ with individual packs nested in regions. Our MixSIAR modelling suggests that Baumann and colleagues^[1]^ may possibly have assumed that the individual ‘niches’ are nested^[29]^ within the whole sample of all Magdalenian canids. This would be permissible^[33]^ “*provided all groups are directly comparable (i*.*e. these groups had access to the exact same set of resources, have similar variances and correlations)*”. This is in contradiction to the conclusion that a close proximity of some of the canids and humans introduced *“a controlled, or at least a restrictive diet*”. As a result, the dietary mix of each ‘trophic niche’ is not reconstructed independently but is calculated taking into account the estimated dietary mix of the whole sample. The authors did not report this, nor did they provide any arguments why this assumption is appropriate. If the three niches were indeed ‘trophic niches’ this nesting could be inappropriate, because the three niches would then occupy separate and independent ecological spaces. We want to stress that we can here only surmise the MixSIAR model settings and assumptions. If Baumann and colleagues^[1]^ used different settings, they should have reported them.

To illustrate the importance of assumptions on model outcomes, we ran separate diet reconstructions for each individual ‘trophic niche’ without any nesting. To this purpose we adapted the example script “mixsiar_script_palmyra.R”^[29]^, which uses the raw data of all source specimens. As no reason for, or information on dependencies between the ‘trophic niches’ is provided by Baumann and colleagues^1^, this would be a reasonable default assumption. In all our simulations we used an ‘uninformative prior’ and we presume Baumann and colleagues^1^ also did so, although they do not mention any prior at all. We provided MixSIAR with the fox TEFs reported by Baumann and colleagues^[1]^. The results of this dietary reconstruction differ significantly from those reported by Baumann and colleagues^[1]^ (Fig. 3c). The dietary preferences are now much less distinct and almost the same for ‘trophic niches’ B and C. If one applies the same non-hierarchical dietary reconstruction on clusters based on separate TEFs for foxes and large canids (Figs. 1d and 2c), results change again (Fig. 3d). Now the modified ‘dog’s niche’ shows a higher proportion of small game in the diet than the ‘fox’s niche’, contrary to the results presented by Baumann and colleagues^[1]^. The clear differences in dietary mixes for the three ‘trophic niches’ presented by Baumann and colleagues^[1]^ result from the unstated model assumptions. Without these, the results are less convincing. We do not want to claim that any of our dietary reconstructions is correct. This exercise merely illustrates that the modelled dietary mix, of course, depends on the model settings and assumptions, which should have been, but were not stated and critically discussed.

Baumann and colleagues^[1]^ write in their SI that *“To get a robust statistical analysis, we set the MCMC (Markov Chain Monte Carlo81) chain length to 1,000,000 with a burn-in of 500,000 in 3 chains”*. They also report the statistical tests for convergence of the model. This may give the impression of statistically responsible science but distracts from the fact that the most basic input parameters and assumptions for their modelling were not reported. It obfuscates that the results may be overly optimistic and biased to build a case for a far-reaching “*proposal for the origin of dogs*” as the paper title purports.

We here do not want to determine an alternative or even better estimate of the dietary mixes of the three ‘trophic niches’. Firstly, we already argued that the ‘trophic niches’ may be spurious, and one may doubt whether TEFs for foxes should be applied to large canids. Secondly, a dietary reconstruction is only meaningful when all major prey sources are known. That most foxes plot outside the convex hull of the three source groups indicates that a low δ^13^C source is probably missing. In a parallel study, Baumann and colleagues^[34]^ carried out similar dietary reconstructions for pre-LGM canids where they included voles and lemmings as sources. These were found to have a wide range of isotopic values in the low δ^13^C and δ^15^N range, but also almost reaching the high δ^15^N values of mammoths. Consumption of voles or lemmings (not unlikely in the Magdalenian) could possibly explain the isotopic values of canids in ‘trophic niche’ C, but this is not mentioned at all. What we here want to point out is that dietary reconstructions may change dramatically depending on the input and assumptions (Fig. 3), including the possible dietary sources^[30]^.

### The taxonomic status of the Gnirshöhle specimens

We finally come to possibly the most critical issue with the study of Baumann and colleagues^[1]^: are the Gnirshöhe specimens dogs or wolves? Although the study uses three specimens from Kesslerloch that show indications of dog domestication^[35]^, it heavily relies on the specimens from the Gnirshöhle site, as is evident from the title of the paper. The Gnirshöhle specimens are labelled *‘large canids’* (*Canis* sp.) as no unequivocal evidence has been published to assign them to either wolves or dogs. The mitochondrial DNA provided no indication of their taxonomic status as the Gnirshöhle specimens are quite diverse and do not systematically cluster or correlate with any wolf or dog lineage. The authors write *‘The phylogenetic arrangement [*…*] did not reveal any clear chronological or spatial differentiation of our six Magdalenian samples compared to the assemblage of ancient and modern canids’*.

In the main text Baumann and colleagues^[1]^ discuss the morphological and metric analysis of only one single Gnirshöhle specimen: the incomplete mandible GN-999. The length of the first lower molar (CLM_1_) is plotted against the length of the tooth row (ALP_1_M_3_) in their Fig. 2. The choice of CLM_1_ as a metric is curious as their Supplementary Information (SI2) states *‘conclusions from measurements of single teeth are not appropriate to classify wolf or dog’*. The use of this metric is therefore misleading in a graph (their Fig. 2) that is purportedly used to classify wolf or dog. The other metric (ALP_1_M_3_) puts GN-999 at the edge of the size range for various wolf (large) and dog (small) groups. The authors write: *‘from the archaeozoological perspective, neither the metrics nor tooth crowding was sufficient to differentiate whether the canids of Gnirshöhle were dogs or wolves’*. If, however, the ALP_1_M_3_-metric would have been a few mm larger, the sample would be clearly in the field for wolves.

Although the assignment of GN-999 to either dog or wolf was unsuccessful, Baumann and collleagues^[1]^ afterwards tacitly assume that all the Gnirshöhle specimens are dogs, as the paper and title state they are crucial to determine the origin of dogs. The ‘trophic niche’ B is labelled with a dog pictogram to strengthen the suggested association of the niche with dogs. This is curious as the metrics of another Gnirshöhle specimen, GN-192, *“places this specimen into the range of Pleistocene and modern wolves”* (their Suppl. Notes 2). The same holds for specimen HF-530 but that specimen was accordingly assigned to the ‘wolves’ of ‘trophic niche’ A. Without any further information, there seems to be no reason to assign the Gnirshöhle specimens to dogs instead of wolves. It seems that the Gnirshöhle specimens were actually assigned to ‘dogs’ or ‘wolves’ depending on their isotopic signature, which would turn the paper into a circular argument.

Importantly, possible evidence that GN-999 (the only Gnirshöhle specimen that was subjected to a morphometric analysis in the main text) may rather be a subadult wolf than a dog was provided, well before submission of the paper, to one of the authors who is listed as supervisor of the study. The pulp chamber of canid teeth fills from the outside inwards^[36,37]^. The CT-scan that was provided (Fig. 4) shows that the pulp cavity infilling ratio of the lower M1 of GN-999 is a low 31% for the mesial, and 34% for the distal root, indicating that GN-999 died soon after full dentition eruption at the age of about six months, well before reaching adult size at an age of about one year^[37]^. The possible subadult status is not mentioned at all in the paper by Baumann and colleagues^[1]^. It is, however, potentially a crucial piece of information on the question whether the Gnirshöhle specimens were dogs at all, as is suggested in the paper. The fact that M_1_ of GN-999 has erupted fully indicates that it was certainly not a young puppy. However, the evidence that GN-999 was sub-adult and did not yet reach its full adult size should be considered and discussed. Only a few per cent (a few mm) additional growth of the mandible to full adult size would have put GN-999 fully in the wolves’ range in terms of ALP_1_M_3_ and beyond that of known modern and ancient dogs (their Fig. 2). Considering this evidence, it is even less probable that the Gnirshöhle specimens would have been dogs. However, by excluding this piece of information, Baumann and colleagues^[1]^ could effectively maintain their tacit assumption that GN-999 is a dog, and then, by inference, all Gnirshöhle specimens in ‘niche B’. The probable subadult status of GN-999 makes the assignment to wolves the most plausible one.

**Figure 4.**
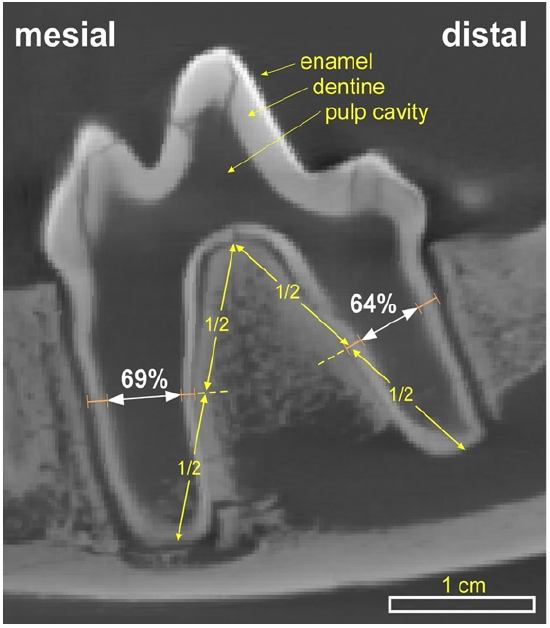
Micro-CT scan of the lower M1 molar of the Gnirshöhle specimen GN-999. It shows a very large pulp cavity (dark) and only a thin layer of dentine (grey). The very early stage of pulp cavity infilling with the pulp cavity width at 69% and 64% indicates that the molar just erupted. This is consistent with the preservation of the thin enamel layer (white) that does not show evidence for abrasion. Measurements of the infilling are according to Nomokonova and colleagues^[37]^.

## Conclusions

The ‘refined proposal for the origin of dogs’ by Baumann and colleagues^[1]^ is based on fundamentally flawed analyses and interpretations.

- The construction of ‘trophic niches’ is wrought with statistical problems (Fig. 1) and based on unsupported assumptions.
- The dietary reconstruction for these ‘trophic niches’ is questionable. The authors refrained from providing the necessary graphs (Fig. 2), model settings and assumptions, nor critically discussed any limitations and alternative options.
- The paper uses tendentious and suggestive graphics and language (such as naming ‘trophic cluster’ B ‘dogs’ with a pictogram of a dog) and provides distracting detail in places (e.g., convergence statistics of MixSIAR modelling), while not reporting critical information, such as the settings of the MixSIAR modelling.
- The line of argument heavily relies on the Gnirshöhle specimens being dogs rather than wolves, but the paper does not provide evidence that these specimens are indeed dogs. Available CT-data that could potentially indicate that the one and only reference Gnirshöhle specimen, GN-999, assigned to dogs, is instead a wolf was not reported.

Finally, we wish to stress that we do not challenge the general validity of (Bayesian) dietary modelling nor the concept of trophic niches, provided rigorous statistics are applied and all assumptions and model settings are clearly reported and critically discussed.

## Data Availability

All data used in this study are derived from published material as referenced in the text. Data from Baumann and colleagues^[1]^ is also listed in the Supplementary Information.

## Supporting information

Method and data for MixSIAR models

## Acknowledgements

C.C.B. thanks the German Research Foundation (DFG) for financial support through grant nr. BA643/1-1. We thank Christoph Wißing for a critical review of the manuscript.

## Author Contributions

P.D.B conceived the study and was the lead author. C.C.B. provided data for the morphometrics. P.D.B and L.P. carried out the MixSIAR modelling. All authors contributed to and reviewed the text and figures.

