## Supplementary material for "Dietary reconstructions of Magdalenian canids from SW-Germany do not indicate that the area was a centre of early European wolf domestication": Method and data for MixSIAR models

We used the data of Baumann and colleagues[1] to run dietary mix reconstructions with the Bayesian model MixSIAR[2]. MixSIAR uses three sets of input data (.cvs files): one for the fraction factors or TEFs, one for the isotope data of the consumers, and one for those of the sources. We here provide the R-scripts for the three reconstructions shown in Fig 3b-d together with the input data that were used for each. We do not provide these as separate files, but as text blocks that can easily be copied and pasted into the appropriate .R or .cvs files that are both essentially text files. We ran the simulations under MacOS 12.6, using RStudio 2022.07.1. We set a Markov Chain Monte Carlo chain length to 1,000,000 with a burn-in of 500,000 in 3 chains.

For the nested or hierarchical dietary mixing model shown in Fig. 3b we used the following R-script:

```
# Script based on mixsiar_script_wolves.R
# Brian Stock
# March 8, 2016
# Script file to run palmyra example without GUI
# Palmyra example ("Taxa" = fixed effect, "raw" source data)
# Remember to first set the working directory with the three input files
mixsiar.dir<- find.package("MixSIAR")
library(MixSIAR)
# Set the file names of the three input files
mix.filename <- "canids_consumer.csv"
source.filename <- "canids_sources.csv"
discr.filename <- "canids_discrimination.csv"
# Load the mixture/consumer data. Here clusters are nested in All_canids
 mix <- load_mix_data(filename=mix.filename,
                      iso_names=c("d13C","d15N"),
                      factors=c("All_canids","Cluster"),
                      fac_random=c(TRUE,TRUE),
                      fac_nested=c(FALSE,TRUE), ## original BC False, True
                      cont_effects=NULL)
# Load the source data
source <- load_source_data(filename=source.filename,
    source_factors="All_canids",
                       conc_dep=FALSE, data_type="means", mix)
# Load the discrimination/TDF data
discr <- load_discr_data(filename=discr.filename, mix)
# Make isospace plot, etc:
plot_data(filename="isospace_plot", plot_save_pdf=FALSE, plot_save_png=TRUE,
    mix, source, discr)
calc_area(source=source,mix=mix,discr=discr)
plot_prior(alpha.prior=1,source)
# Write the JAGS model file
model_filename <- "MixSIAR_model.txt"   # Name of the JAGS model file
resid_err <- TRUE # normal TRUE
process_err <- TRUE # normal TRUE
write_JAGS_model(model_filename, resid_err, process_err, mix, source)
# MCMC run options:
# run <- "test" # chainLength=1000, burn=500, thin=1, chains=3, calcDIC=TRUE
# run <- "very short" # chainLength=10000, burn=5000, thin=5, chains=3,
    calcDIC=TRUE
# run <- "short" # chainLength=50000, burn=25000, thin=25, chains=3,
    calcDIC=TRUE
# run <- "normal" # chainLength=100000, burn=50000, thin=50, chains=3,
    calcDIC=TRUE
```

```
# run <- "long" # chainLength=300000, burn=200000, thin=100, chains=3,
    calcDIC=TRUE
# run <- "very long" # chainLength=1000000, burn=500000, thin=500, chains=3,
    calcDIC=TRUE
# run <- "extreme" # chainLength=3000000, burn=1500000, thin=500, chains=3,
    calcDIC=TRUE
# After a test run works, choose the appropriate MCMC to run the model
jags.1 <- run_model(run="test", mix, source, discr, model_filename, alpha.prior
    = 1)
# Set output options
output_options <- list(summary_save = TRUE,
                       summary_name = "summary_statistics",
                       sup_post = TRUE,
                       plot_post_save_pdf = TRUE,
                       plot_post_name = "posterior_density",
                       sup_pairs = FALSE,
                       plot_pairs_save_pdf = TRUE,
                       plot_pairs_name = "pairs_plot",
                       sup_xy = TRUE,
                       plot_xy_save_pdf = TRUE,
                       plot_xy_name = "xy_plot",
                       gelman = TRUE,
                       heidel = TRUE,
                       geweke = TRUE,
                       diag_save = TRUE,
                       diag_name = "diagnostics",
                       indiv_effect = FALSE,
                       plot_post_save_png = TRUE,
                       plot_pairs_save_png = TRUE,
                       plot_xy_save_png = TRUE)
# Get diagnostics, summary statistics, and posterior plots
 df.stats <- output_stats(jags.1, mix, source, output_options)
 df.diag <- output_diagnostics(jags.1, mix, source, output_options)
 g.post <- output_posteriors(jags.1, mix, source, output_options)
 output_JAGS(jags.1, mix, source, output_options)
```

The isotope values of the three dietary sources were provided as means and their standard deviation. The three input data files are:

The file canids_consumer.cvs:

```
d13C,    d15N,   Region,  Pack
-19.5,   9.1,    1,       1
-19.4,   7.4,    1,       1
-20.1,   7.1,    1,       1
-20.0,   7.9,    1,       1
-19.7,   8.2,    1,       1
-19.2,   6.1,    1,       2
-19.8,   5.6,    1,       2
-19.9,   5.3,    1,       2
-19.4,   6.1,    1,       2
-19.5,   5.6,    1,       2
-19.9,   5.9,    1,       2
-19.0,   5.7,    1,       2
-19.1,   6.4,    1,       2
-19.2,   5.9,    1,       2
-19.6,   4.7,    1,       2
-20.3,   5.8,    1,       3
-20.6,   5.3,    1,       3
-20.3,   5.6,    1,       3
-20.6,   5.1,    1,       3
-20.6,   4.5,    1,       3
```

The file canids_discrimination.cvs with TEFs for foxes[3]:

```
,              Meand13C,   SDd13C,  Meand15N,   SDd15N
megafauna,     1.1,        1.1,     3.2,        1.8
ungulates,     1.1,        1.1,     3.2,        1.8
small_game,    1.1,        1.1,     3.2,        1.8
```

The file candids_sources.cvs:

```
,               Region,   Meand13C,   SDd13C,   Meand15N,   SDd15N,   n
megafauna,      1,        -20.97,     0.42,     5.87,       0.76,     3
ungulates,      1,        -19.85,     0.42,     2.2,        0.77,     69
small_game,     1,        -20.52,     0.39,     1.24,       0.72,     21
```

The R-script for the second set of mixing models of Fig. 3c and 3d is given below. Here we ran non-hierarchical mixing models on the three 'clusters' of consumers separately. Sources were provided as raw data, i.e., the isotope values of the individuals.

```
# Script based on:
# Brian Stock
# March 8, 2016
# Script file to run palmyra example without GUI
# Palmyra example ("Taxa" = fixed effect, "raw" source data)
# Remember to first set the working directory with the three input files
mixsiar.dir<- find.package("MixSIAR")
library(MixSIAR)
# Set the file names of the three input files
mix.filename <- "canids_consumer.csv"
source.filename <- "canids_sources.csv"
discr.filename <- "canids_discrimination.csv"
# Load the mixture/consumer data
mix <- load_mix_data(filename=mix.filename,
                         iso_names=c("d13C","d15N"),
                         factors="Taxa",
                         fac_random=FALSE,
                         fac_nested=tRUE, #standard FALSE
                         cont_effects=NULL)
# Load source data
source <- load_source_data(filename=source.filename,
                                  source_factors=NULL,
                                  conc_dep=FALSE,
                                  data_type="raw",
                                  mix)
# Load discrimination/TDF data
discr <- load_discr_data(filename=discr.filename, mix)
# Make isospace plot, etc
plot_data(filename="isospace_plot",
      plot_save_pdf=TRUE,
      plot_save_png=FALSE,
      mix,source,discr)
# Calculate standardized convex hull area
if(mix$n.iso==2) calc_area(source=source,mix=mix,discr=discr)
# Plot your prior
 plot_prior(alpha.prior=1,source)
# Define model structure and write JAGS model file
model_filename <- "MixSIAR_model.txt"
resid_err <- TRUE # FALSE if only one mix point
process_err <- TRUE
write_JAGS_model(model_filename, resid_err, process_err, mix, source)
# MCMC run options:
# run <- "test"       # chainLength=1000, burn=500, thin=1, chains=3,
    calcDIC=TRUE
# run <- "very short" # chainLength=10000, burn=5000, thin=5, chains=3,
    calcDIC=TRUE
# run <- "short"      # chainLength=50000, burn=25000, thin=25, chains=3,
    calcDIC=TRUE
# run <- "normal"     # chainLength=100000, burn=50000, thin=50, chains=3,
    calcDIC=TRUE
# run <- "long"       # chainLength=300000, burn=200000, thin=100, chains=3,
    calcDIC=TRUE
# run <- "very long"  # chainLength=1000000, burn=500000, thin=500, chains=3,
    calcDIC=TRUE
# run <- "extreme"    # chainLength=3000000, burn=1500000, thin=500, chains=3,
    calcDIC=TRUE
# After a test run works, choose the appropriate MCMC to run the model
```

```
jags.1 <- run_model(run="test", mix, source, discr, model_filename,
    alpha.prior=1)
output_options <- list(summary_save = TRUE,
                       summary_name = "summary_statistics",
                       sup_post = TRUE,
                       plot_post_save_pdf = TRUE,
                       plot_post_name = "posterior_density",
                       sup_pairs = FALSE,
                       plot_pairs_save_pdf = TRUE,
                       plot_pairs_name = "pairs_plot",
                       sup_xy = TRUE,
                       plot_xy_save_pdf = TRUE,
                       plot_xy_name = "xy_plot",
                       gelman = TRUE,
                       heidel = FALSE,
                       geweke = TRUE,
                       diag_save = TRUE,
                       diag_name = "diagnostics",
                       indiv_effect = FALSE,
                       plot_post_save_png = FALSE,
                       plot_pairs_save_png = FALSE,
                       plot_xy_save_png = TRUE)
# Get diagnostics, summary statistics, and posterior plots
df.stats <- output_stats(jags.1, mix, source, output_options)
df.diag <- output_diagnostics(jags.1, mix, source, output_options)
g.post <- output_posteriors(jags.1, mix, source, output_options)
output_JAGS(jags.1, mix, source, output_options)
```

For Fig. 3c, all canids were given the same TEF for foxes[3]:

The file canids_consumer.cvs (niche A):

```
Taxa,     d13C,    d15N
Niche_A,  -19.5,   9.1
Niche_A,  -19.4,   7.4
Niche_A,  -20.1,   7.1
Niche_A,  -20,     7.9
Niche_A,  -19.7,   8.2
```

The file canids_consumer.cvs (niche B):

```
Taxa,     d13C,    d15N
Niche_B,  -19.2,   6.1
Niche_B,  -19.8,   5.6
Niche_B,  -19.9,   5.3
Niche_B,  -19.4,   6.1
Niche_B,  -19.5,   5.6
Niche_B,  -19.9,   5.9
Niche_B,  -19,     5.7
Niche_B,  -19.1,   6.4
Niche_B,  -19.2,   5.9
Niche_B,  -19.6,   4.7
```

The file canids_consumer.cvs (niche C):

```
Taxa,     d13C,    d15N
Niche_C,  -20.3,   5.8
Niche_C,  -20.6,   5.3
Niche_C,  -20.3,   5.6
Niche_C,  -20.6,   5.1
Niche_C,  -20.6,   4.5
```

The file canids_discrimination.cvs with TEFs for foxes[3]:

```
,              Meand13C,   SDd13C,  Meand15N,   SDd15N
megafauna,     1.1,        1.1,     3.2,        1.8
ungulates,     1.1,        1.1,     3.2,        1.8
small_game,    1.1,        1.1,     3.2,        1.8
```

The file canids_sources.cvs:

```
,              d13C,     d15N
megafauna,     -21.3,    5
```

```
megafauna,     -20.5,     6.4
megafauna,     -21.1,     6.2
small_game,    -20.8,     1.9
small_game,    -20.8,     0.9
small_game,    -20.2,     0.7
small_game,    -20.8,     0.5
small_game,    -21.4,     0.2
small_game,    -20.6,     0.3
small_game,    -20.4,     2.3
small_game,    -19.7,     1.3
small_game,    -20.2,     0.4
small_game,    -20.4,     0.6
small_game,    -20.5,     1.1
small_game,    -20.2,     0.5
small_game,    -20.1,     1.9
small_game,    -20.4,     1
small_game,    -20.3,     1.6
small_game,    -20.2,     0.9
small_game,    -20.9,     1.5
small_game,    -20.9,     1.7
small_game,    -20.3,     2
small_game,    -21,       2.5
small_game,    -20.8,     2.2
ungulates,     -19.8,     2.2
ungulates,     -20,       2.4
ungulates,     -20.3,     2.3
ungulates,     -20.2,     3
ungulates,     -19.8,     2.9
ungulates,     -20.7,     2.2
ungulates,     -20.1,     2.9
ungulates,     -19.8,     1.6
ungulates,     -20.1,     1.5
ungulates,     -20.4,     1.3
ungulates,     -20.6,     2.1
ungulates,     -20.6,     1.3
ungulates,     -20.4,     1.6
ungulates,     -20.2,     2.3
ungulates,     -20.3,     0.6
ungulates,     -20,       1.7
ungulates,     -20.8,     2.1
ungulates,     -19.9,     0.8
ungulates,     -20.4,     2
ungulates,     -20.6,     2.4
ungulates,     -19.5,     3.7
ungulates,     -19.5,     2.4
ungulates,     -19.1,     2.8
ungulates,     -19.1,     3.9
ungulates,     -19.3,     1.7
ungulates,     -19.7,     2.7
ungulates,     -19.5,     1.3
ungulates,     -20.3,     2.1
ungulates,     -20.2,     2
ungulates,     -19.7,     1.7
ungulates,     -19.8,     2.1
ungulates,     -19.3,     2.2
ungulates,     -20.2,     2.1
ungulates,     -19.4,     0.1
ungulates,     -19.6,     2
ungulates,     -19.4,     1
ungulates,     -19.1,     2.4
ungulates,     -19.9,     2.6
ungulates,     -19.8,     2.7
ungulates,     -20.3,     2.9
ungulates,     -19.9,     2.8
ungulates,     -19.8,     2.4
ungulates,     -19.6,     2.1
ungulates,     -19.9,     2.3
ungulates,     -20,       1.2
```

```
ungulates,     -19.9,     1.2
ungulates,     -19.4,     2.4
ungulates,     -20.2,     2.6
ungulates,     -19.1,     2.3
ungulates,     -19.7,     2.1
ungulates,     -19.7,     2.5
ungulates,     -19.7,     2.5
ungulates,     -19.7,     1.4
ungulates,     -20,       3.3
ungulates,     -19.7,     2.4
ungulates,     -19.9,     2
ungulates,     -19.5,     2.6
ungulates,     -19.6,     2.4
ungulates,     -19.3,     2.3
ungulates,     -19.1,     1.9
ungulates,     -19.4,     2.9
ungulates,     -19.8,     2.5
ungulates,     -19.7,     2.8
ungulates,     -20.1,     1.8
ungulates,     -19.4,     1.9
ungulates,     -19.5,     2.2
ungulates,     -20.3,     5.3
ungulates,     -19.9,     2.1
```

The third set of models (Fig. 3d) is for individual TEFs for large canids[4] and for foxes[3], resulting in modified groups (Figs. 1d and 2c). Consumer data in the canids_consumer.cvs files are now corrected for the different TEFs, and the canids_discrimination.cvs file now only contains the one standard deviation uncertainty in the TEFs, for which the (smaller) value for the wolves' TEF[4] was taken. This set of models used the same source input as above.

The file canids_discrimination.cvs with individual TEFs for large canids[4] and for foxes[3]:

```
,      Meand13C,     SDd13C,       Meand15N,     SDd15N
megafauna,    0,     0.6,   0,     0.7
ungulates,    0,     0.6,   0,     0.7
small_game,   0,     0.6,   0,     0.7
```

The file canids_consumer.cvs (modified niche A):

```
Taxa,         d13C,  d15N
NewNiche_A,  -20.8, 4.5
NewNiche_A,  -20.8, 5
```

The file canids_consumer.cvs (modified niche B):

```
Taxa,         d13C,  d15N
NewNiche_B,  -20.7, 2.8
NewNiche_B,  -20.5, 1.5
NewNiche_B,  -20.7, 1.5
NewNiche_B,  -20.8, 1
NewNiche_B,  -20.3, 1.1
NewNiche_B,  -20.4, 1.8
NewNiche_B,  -20.5, 1.3
NewNiche_B,  -20.7, 1.5
```

The file canids_consumer.cvs (modified niche C):

```
Taxa,         d13C,  d15N
NewNiche_C,  -21.4, 2.5
NewNiche_C,  -21.3, 3.3
NewNiche_C,  -21.1, 1
NewNiche_C,  -21.2, 0.7
NewNiche_C,  -21.2, 1.3
NewNiche_C,  -21.6, 1.2
NewNiche_C,  -21.7, 2.1
NewNiche_C,  -21.4, 2.4
NewNiche_C,  -21.7, 1.9
```

```
NewNiche_C,  -21.7, 1.3
```
